# Narcolepsy with cataplexy is caused by epigenetic silencing of hypocretin neurons

**DOI:** 10.1101/2021.09.21.461046

**Authors:** Ali Seifinejad, Almar Neiteler, Sha Li, Corinne Pfister, Rolf Fronczek, Ling Shan, Lee A. Garrett-Sinha, David Frieser, Makoto Honda, Yoan Arribat, Dogan Grepper, Francesca Amati, Marie Picot, Christian Iseli, Nicolas Chartrel, Gert J. Lammers, Roland Liblau, Anne Vassalli, Mehdi Tafti

## Abstract

Narcolepsy with cataplexy is a chronic sleep disorder characterized by hypocretin deficiency. The condition is believed to result from autoimmune destruction of hypocretin (HCRT) neurons, although direct evidence is lacking and mere *Hcrt* gene inactivation causes full-blown narcolepsy in mice. Here we show that the expression of another hypothalamic neuropeptide, QRFP, is lost in mouse models with HCRT cell-ablation, but tends to be even increased in *Hcrt* gene knockout mice, suggesting that QRFP expression can be used as a proxy for the presence or absence of HCRT neurons. Similar to *Hcrt* knockout mice, narcolepsy patients show intact hypothalamic *QRFP* expression, and cerebrospinal fluid levels of QRFP peptide are increased, suggesting their HCRT neurons are intact. We show that the human *HCRT* gene promoter is methylation-sensitive *in vitro*, and is hypermethylated in the hypothalamus of patients selectively at a putative PAX5:ETS1 binding site within the proximal *HCRT* promoter. Ets1-KO mice display downregulated *Hcrt* expression, while *pax5-ets1* knockdown in zebrafish causes decreased *hcrt* expression, decreased activity and sleep fragmentation, similar to narcolepsy patients. Our results suggest that HCRT neurons are alive, but epigenetically silenced, in the hypothalamus of narcolepsy patients, opening the possibility to reverse or cure narcolepsy.

---

Narcolepsy with cataplexy (also called narcolepsy type 1 or NT1, hereafter called narcolepsy) is a sleep disorder characterized by excessive daytime sleepiness and cataplexy (loss of skeletal muscle tone triggered by strong emotions)^1^. The best biological marker of the disease is hypocretin (HCRT) deficiency (cerebrospinal fluid [CSF] HCRT-1 levels <110 pg/ml)^2^. Post-mortem examination of the hypothalamic region of narcolepsy patients indicated a near absence of HCRT-producing neurons^3,4^. Together with strong associations with the *HLA-DQB1*06:02* allele^5^, a T cell receptor alpha gene polymorphism^6^, and the acute increase in childhood narcolepsy following the 2009-2010 H1N1 vaccination^7^, it is widely, if not unanimously, accepted that an immune or autoimmune process leads to HCRT neuronal loss. Although autoreactive T cells against various antigens expressed by HCRT neurons were recently reported^8,9^, a direct evidence of HCRT neuronal damage (cell death, lateral hypothalamic inflammation or T cell infiltration) is still lacking.

HCRT deficiency can be speculated to result from either neuronal death or lack of HCRT production. Evidence from patients with secondary narcolepsy^10-13^ indicates that CSF HCRT-1 deficiency can be temporary and normalize following successful treatment, suggesting that hypothalamic inflammatory reactions may lead to reversible HCRT deficiency. Interestingly, a 28-year-old female patient diagnosed with narcolepsy with cataplexy and undetectable CSF HCRT-1, was treated with intravenous immunoglobulins starting only 15 days after her first cataplexy attack, after which her CSF HCRT-1 level normalized, and her symptoms improved^14^. Also, up to 48% of patients with traumatic brain injury exhibit HCRT deficiency, which recovers within 6 months^15^. Moreover, altered CSF HCRT-1 levels, and even HCRT deficiency, were reported during hypersomnia episodes in patients with Kleine-Levin syndrome, suggesting changes in expression levels without permanent cell loss^16^. Overall, these data suggest that *HCRT* gene expression can be finely tuned and is sensitive to local and external environmental changes.

If HCRT deficiency is not caused by cellular destruction, then either loss-of-function mutations or epigenetic silencing are involved. Extensive genetic analyses of narcolepsy have failed to uncover causative gene mutations, with the exception of a few cases^3,17,18^, and the disease is thought to be primarily sporadic. Epigenome-wide association studies have revealed sites of differential DNA methylation in blood and hypothalamic samples of narcolepsy patients^19,20^, but no site-specific hypermethylation associated with HCRT deficiency was found.

## Preserved brain QRFP expression in narcolepsy

In a recent study we used RNA sequencing to identify differentially expressed genes in the hypothalamus, cortex and brainstem from HCRT-neuron-ablated (Hcrt-ataxin transgenic mice), Hcrt-KO and littermate control mice^21^. A single gene called *Qrfp* showed a striking hypothalamus-specific 50% decrease in expression, in the hypothalamus of Hcrt-ataxin mice, while levels were unchanged in Hcrt-KO mice. To confirm this RNA-seq finding in another HCRT-neuron-ablated mouse model, we quantified *Qrfp* transcript in the hypothalamus of mice harboring a *HCRT* promoter-driven diphtheria toxin A transgene, another HCRT-neuron-ablated mouse model (Hcrt-DTA)^22^. *Qrfp* expression was found to be profoundly decreased (up to 10-fold) in these mice (Fig. 1a). No expression change was however observed for *Pmch*, the MCH peptide-encoding gene which is also expressed in the lateral hypothalamus in cells intermingled with HCRT and QRFP-producing neurons (Fig. 1a). These findings therefore strongly support the use of *Qrfp* expression level as a proxy for HCRT neuronal loss.

**Fig. 1.**
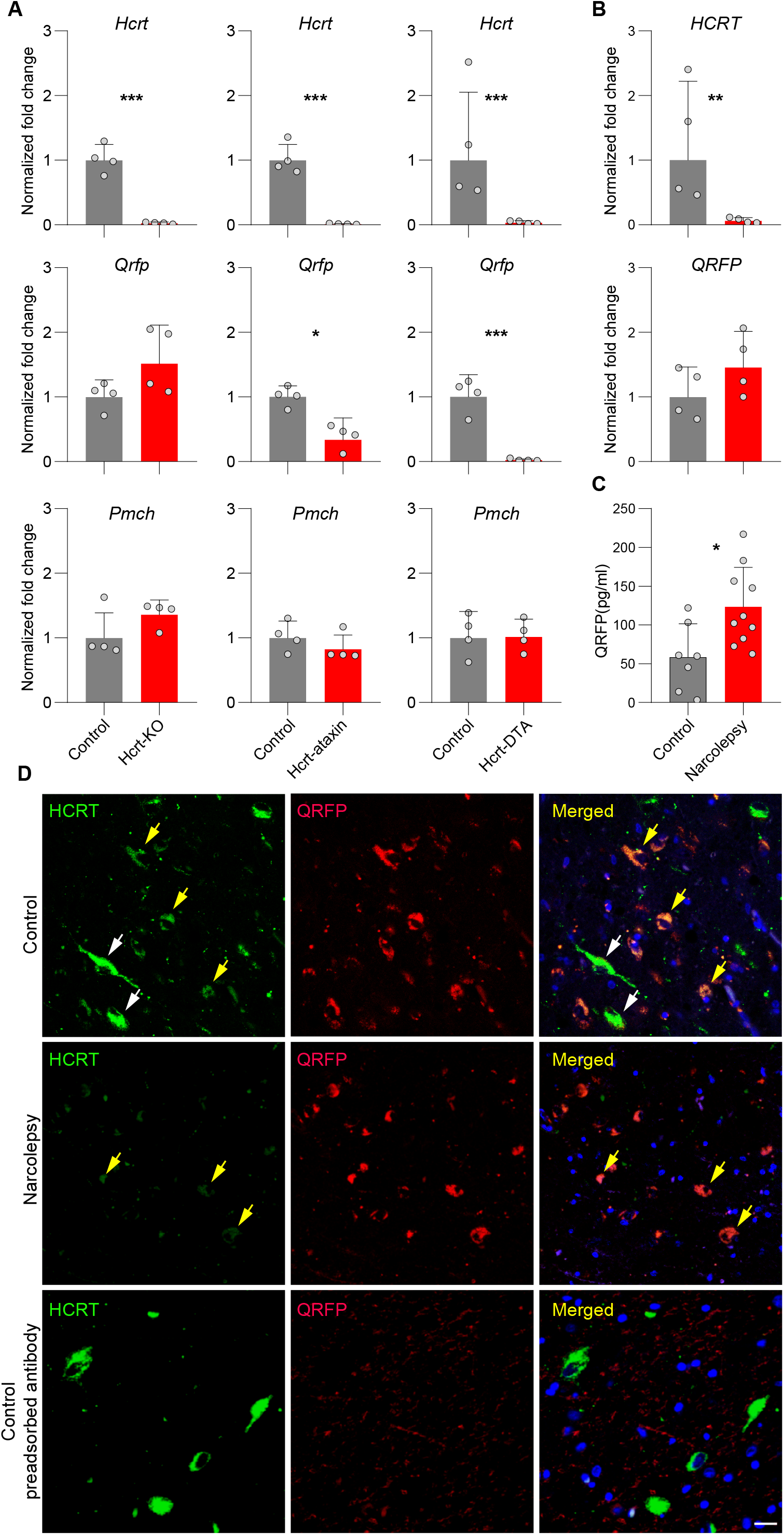
*Qrfp* expression is preserved in the hypothalamus of narcolepsy patients. **a**, Expression levels of *Hcrt, Qrfp*, and *Pmch* in the brain of Hcrt-KO mice and two Hcrt neuron-ablated transgenic lines, Hcrt-ataxin and Hcrt-DTA mice(N=4 in each group). *Hcrt* expression is dramatically reduced in all three narcolepsy mouse models as compared to their respective wild-type controls. *Qrfp* expression is unchanged (or tends to increase) in Hcrt-KO mice, while significantly decreased in both neuron-ablated models. *Pmch* is unchanged in all three mouse models. * *P* < 0.05, ** *P* < 0.01, and *** *P* < 0.001, *t*-tests on log transformed normalized fold change of KO models compared to their controls, N=4. Bars represent geometric means + geometric SD. **b**, *HCRT* expression is also dramatically decreased in the hypothalamus of narcolepsy patients, as expected, while *QRFP* expression tends to be increased, similar to Hcrt-KO mice.. * *P* < 0.05, *t*-test on log transformed normalized fold change of narcolepsy patients compared to controls, N=4. Bars represent geometric means + geometric SD. **c**, QRFP quantification in the CSF by RIA of narcolepsy (N=12) and control subjects (N=7), evidence increased levels in narcolepsy patients. * *P* < 0.05, *t*-test. Bars represent means + SD. **d**, Immunofluorescence staining of postmortem human hypothalamic sections. HCRT, QRFP and merged immunofluorescence in a normal subject (upper panels), a narcolepsy patient (middle panels), or a control subject after pre-adsorption of the antiserum with QRFP peptide. Note that neurons with low HCRT immunoreactivity (yellow arrows) appear QRFP-positive, while neurons with high HCRT expression (white arrows) do not. In the patient, again neurons with low intensity staining for HCRT show strong QRFP staining, and there is no apparent difference in the number of QRFP-positive neurons compared to the control subject. Lower panels of a control subject with QRFP peptide-preadsorbed QRFP antibody indicate the antibody specificity (scale bars: 20 μm).

To determine whether *QRFP* is decreased in the hypothalamus of narcolepsy patients as it is in HCRT neuron-ablated mouse models, RNA was extracted from the hypothalamic region of 4 narcolepsy and 4 control fresh frozen post-mortem brains^23^. Strikingly, *QRFP* expression was not lost in patients’ hypothalamus (Fig. 1b). Not only the expression of *QRFP* was preserved, but it showed a very similar tendency to increase in narcolepsy patients (45% higher than controls), as in Hcrt-KO mice (51% higher than wild-type, Fig. 1a). This may suggest a form of transcription compensation. Note that not only *Qrfp* and *Hcrt* cDNAs share 50% sequence similarity, they also are functionally related and their respective 7-transmembrane receptors can heterodimerize^24^). To support this at the protein level, we also measured the level of QRFP in the CSF of 10 narcolepsy patients and 7 control subjects. QRFP was significantly increased by 2 folds in patients (Fig. 1c), suggesting that CSF QRFP measures may be used as a new biomarker of narcolepsy. Altogether, these results suggest that HCRT neurons might be present in narcolepsy brains but the gene is inactive.

There have been conflicting reports on whether *Qrfp* and *Hcrt* are co-expressed in the same neurons. Distinct hypothalamic neuronal populations in adult mice have been reported^25^, while single-cell RNA-sequencing of hypothalamic neurons indicated co-localization, and hierarchical clustering defined by molecular fingerprints have suggested that HCRT- and QRFP-expressing neurons share a similar neuronal lineage^26^. To test if HCRT and QRFP are co-localized in humans, we performed double immunostaining of post-mortem hypothalamic sections. Analysis of hypothalamic sections (N=4) from four control subjects indicated 50.50±18.63% (mean±sd) colocalization of HCRT and QRFP (Fig. 1d). The same analysis in sections (N=4) from 2 narcolepsy patients did not indicate decrease in number of QRFP-positive neurons, relative to controls (patients: 69.50%; controls: 82%). Amongst the rare HCRT-immunoreactive neurons of patients 80.98% were QRFP-immunoreactive positive (Fig. 1d). Interestingly, the intensity of HCRT and QRFP staining was inversely correlated, with neurons with low HCRT staining showing high level of QRFP expression and *vice versa* (Fig. 1d), again suggesting a causal relationship between the expression of *HCRT* and *QRFP*. Recently, Takahashi *et al*., reported the generation of a knock-in mouse line expressing iCre from the endogenous *Qrfp* gene and observed low levels of iCre expression in HCRT immunoreactive neurons, supporting Hcrt/Qrfp co-expression^27^. This strongly suggest that the loss of HCRT neurons could be indexed by the concomitant loss in QRFP expression, and the fact that expression of *QRFP* mRNA and protein is preserved in narcolepsy patients supports the survival of these cells.

### Epigenetic alteration of *HCRT* gene in narcolepsy

In contrast to HCRT neuron-ablated mouse lines, human narcolepsy brains seem to contain HCRT neurons, but as Hcrt-KO mice they do not express the *HCRT* gene. Since the *HCRT* gene coding sequence is unaffected in narcolepsy, lack of expression may result from epigenetic dysregulation. Epigenetic regulation of gene expression can operate in different ways, including DNA methylation, histone modification, chromatin structural change, and microRNAs. DNA methylation and histone modification are major mechanisms for gene silencing, which mainly occur in regulatory regions of genes such as in promoters and enhancers.

To determine if promoter methylation causes silencing of human *HCRT* expression, we first performed a luciferase assay. A 1800 bp proximal fragment of the human *HCRT* promoter, was cloned in a CpG-free luciferase reporter vector (pCpGL). Two different cell lines (HEK 293FT and PC12) were transfected with the intact unmethylated human *HCRT* promoter-luciferase pCpGL construct, or with the same construct which was exposed to the M.Sssl methyltransferase *in vitro*. Methylation of the *HCRT* promoter decreased the expression of luciferase by 60 to 90% (p<0.05) in these cell lines (Fig. 2a), confirming that methylation of the *HCRT* promoter inhibits *HCRT* expression.

**Fig. 2.**
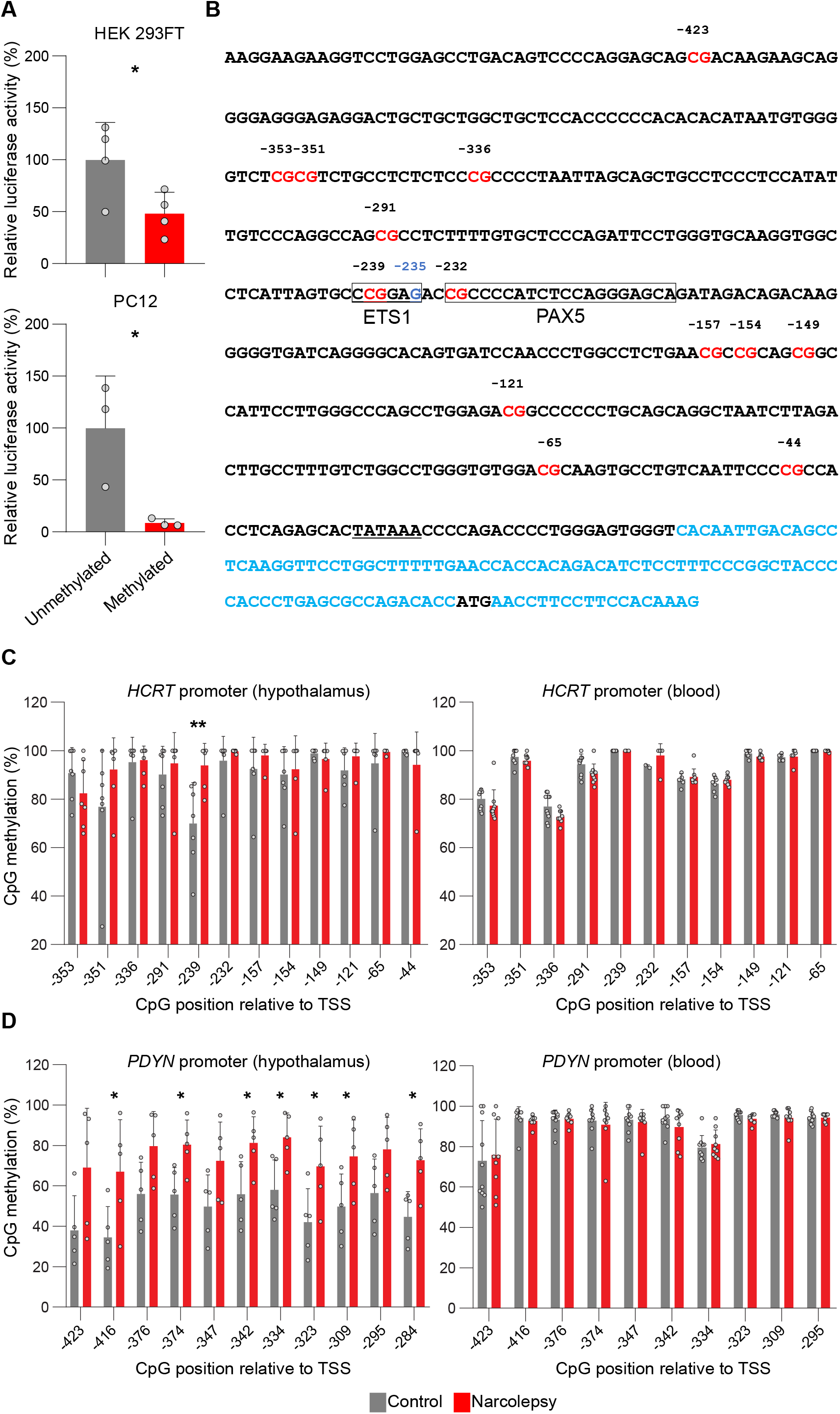
*HCRT* gene promoter is hypermethylated selectively in hypothalamus of narcolepsy patients. **a**, Expression of a human *HCRT* gene promoter-driven luciferase reporter in two different cell lines is suppressed by methylation. Both cultures were transfected with a *HCRT*-promoter-driven luciferase construct either methylated or unmethylated. Values are averages of three wells measured in duplicate over three different transfections and reported as a percentage of the reference unmethylated vector. * *P* < 0.05, *t*-tests of methylated compared to unmethylated, N=4 (HEK 293FT) and N=3 (PC12). Bars represent means + SD. **b**, Sequence of the human *HCRT* promoter. Shown are 500 bp upstream of the transcription start site, together with the first exon (in blue) with 5’ untranslated region (in black). CpGs are highlighted in red. The PAX5:ETS1 binding site is boxed with the G instead of A from the consensus ETS1 sequence in bleu (−235). The TATA box is underlined and the translation start site (ATG) in black. **c**, (left panel) Methylation status at the 12 most proximal CpGs of the *HCRT* promoter (see Figure **2b**) from hypothalamic brain sections in 7 narcolepsy patients and 7 controls. One of the 12 CpG’s is significantly hypermethylated in narcolepsy patients relative to controls (−239 CpG). **c**, (right panel) Methylation status of 11 out of the 12 most proximal CpGs of the *HCRT* promoter from peripheral blood samples of 10 narcolepsy and 10 control subjects (note the hypothalamic and blood samples are not from the same subjects). **d**, (left panel) Methylation status of *PDYN* promoter from brain hypothalamic sections in five narcolepsy and five control subjects showing significant hypermethylation of 8 out of the 11 CpGs relative to controls. **d**, (right panel) Methylation status of *PDYN* promoter from peripheral blood samples of 9 narcolepsy and 10 control subjects. Data for hypothalamic *HCRT* and *PDYN* promoters are from targeted next-generation bisulfite sequencing, while those for blood are from Sanger sequencing (percentage + SD). * *P* < 0.05 and ** *P* < 0.005, *t*-tests between narcolepsy patients and controls. CpG positions are indicated relative to the transcriptional start site (TSS).

To determine whether promoter methylation is involved in the HCRT deficiency of narcolepsy patients, we next evaluated *HCRT* gene promoter methylation in DNA samples extracted from post-mortem lateral hypothalami of seven narcolepsy with cataplexy patients and seven control subjects. The proximal human *HCRT* promoter contains 13 CpGs (Fig. 2b). We applied bisulfite treatment, followed by either of two sequencing methods, to quantify the CpG methylation status. The first method relies on direct Sanger sequencing of PCR amplicons and measures the average methylation of DNA molecules in a tissue. If a differential methylation pattern was observed with this first approach, we performed targeted next-generation bisulfite sequencing. 12 of the 13 CpG sites (all except the first CpG in Fig. 2b) were tested. A higher methylation level was observed in narcolepsy patients as compared to controls at 8 of these 12 CpGs, and one of them was significantly hypermethylated both by Sanger and targeted sequencing (CpG -239, methylation 93.9% in patients vs. 69.93% in controls, p<0.01, Fig. 2c, left panel).

Post-mortem analysis of narcolepsy patients’ brains indicated that immunostaining for two other neuropeptides, PDYN (Dynorphin) and NPTX2 (Narp), which partially colocalize with HCRT, are also lost in the lateral hypothalamus^28,29^. *In silico* analysis of the promoter regions of these two genes revealed that both harbor a CpG island, suggesting that they may also be silenced in narcolepsy. The *NPTX2* promoter contains a very high density CpG island hindering bisulfite sequencing. We therefore focused on the methylation status of the hypothalamic *PDYN* promoter by applying the same methods as for the *HCRT* promoter. All 11 CpGs tested showed higher methylation in patients than in controls, and of them 7 were significantly hypermethylated (N=5 in each group, p<0.05; Fig. 2d, left panel). Therefore, loss of PDYN in narcolepsy might also be due to promoter methylation rather than cell loss.

To determine whether differential methylation in narcolepsy patients compared to controls is specific to HCRT neurons, we tested the methylation status of the *HCRT* receptor 2 gene (*HCRTR2*). *HCRTR2* is expressed in the hypothalamus including the lateral hypothalamus but HCRT neurons do not express *HCRTR2*^30^. The *HCRTR2* promoter contains a CpG island, in which we evaluated 10 CpGs for methylation level. Surprisingly, we found very low methylation (all CpGs had a methylation level below 35%), and no difference between narcolepsy patients and controls (N=4 narcolepsy patients and 6 controls, Supplementary Fig. 1). This suggests that promoter hypermethylation within the lateral hypothalamus of narcolepsy patients may specifically affect HCRT neurons.

As methylation is cell-specific, we tested methylation of the *HCRT* and *PDYN* promoters in DNA samples extracted from the peripheral blood cells of narcolepsy patients and controls. Eleven out of the same 12 CpGs of *HCRT* as tested in the brain samples were also tested by Sanger sequencing. Similar to the brain DNA, methylation at these 11 CpGs varied between 68 and 100% in the blood DNA. We however found that the CpG -239 that was hypermethylated in the hypothalamus of narcolepsy patients compared to controls was methylated at 100% in peripheral blood of all subjects, in patients or controls (N=10 in each group, Fig. 2c, right panel). The fact that the level of methylation is not dramatically different between blood and brain (although the pattern is), indicates that our hypothalamic samples contain heterogeneous populations of cells and the fraction of the relevant HCRT-producing neurons is small, leading to dilution of the signal. The same analysis was also performed for *PDYN* DNA from blood samples, which likewise indicated high methylation and no difference between narcolepsy patients and controls (N=9 narcolepsy patients and 10 controls, Fig. 2d, right panel). We also examined the CpG methylation levels of *QRFP* and found no changes either in the hypothalamus or blood of narcolepsy patients as compared to controls (N=7 narcolepsy patients and 6 controls for the hypothalamus, N=9 narcolepsy and 10 controls for the blood, Supplementary Fig. 2).

### The PAX5:ETS1 complex regulates *HCRT* promoter

The hypermethylated CpG -239 in the *HCRT* promoter is part of a suboptimal ETS1 binding site (5’-CCGGAG-3’), a sequence originally identified in the promoter of the early B-cell-specific mb-1 (*CD79A*) gene, encoding the Ig-alpha protein^31^. ETS1 binding at this site was reported to be repressed by methylation^32^. To determine whether this site in the *HCRT* promoter recruits ETS1, we mutated this binding sequence to create an optimal ETS1 binding site (5’-CCGGAA-3’) in our *HCRT* promoter-driven luciferase reporter plasmids. The mutated construct showed significantly increased luciferase activity (Fig. 3a), thus demonstrating that the identified binding site is important for *HCRT*-driven expression.

**Fig. 3.**
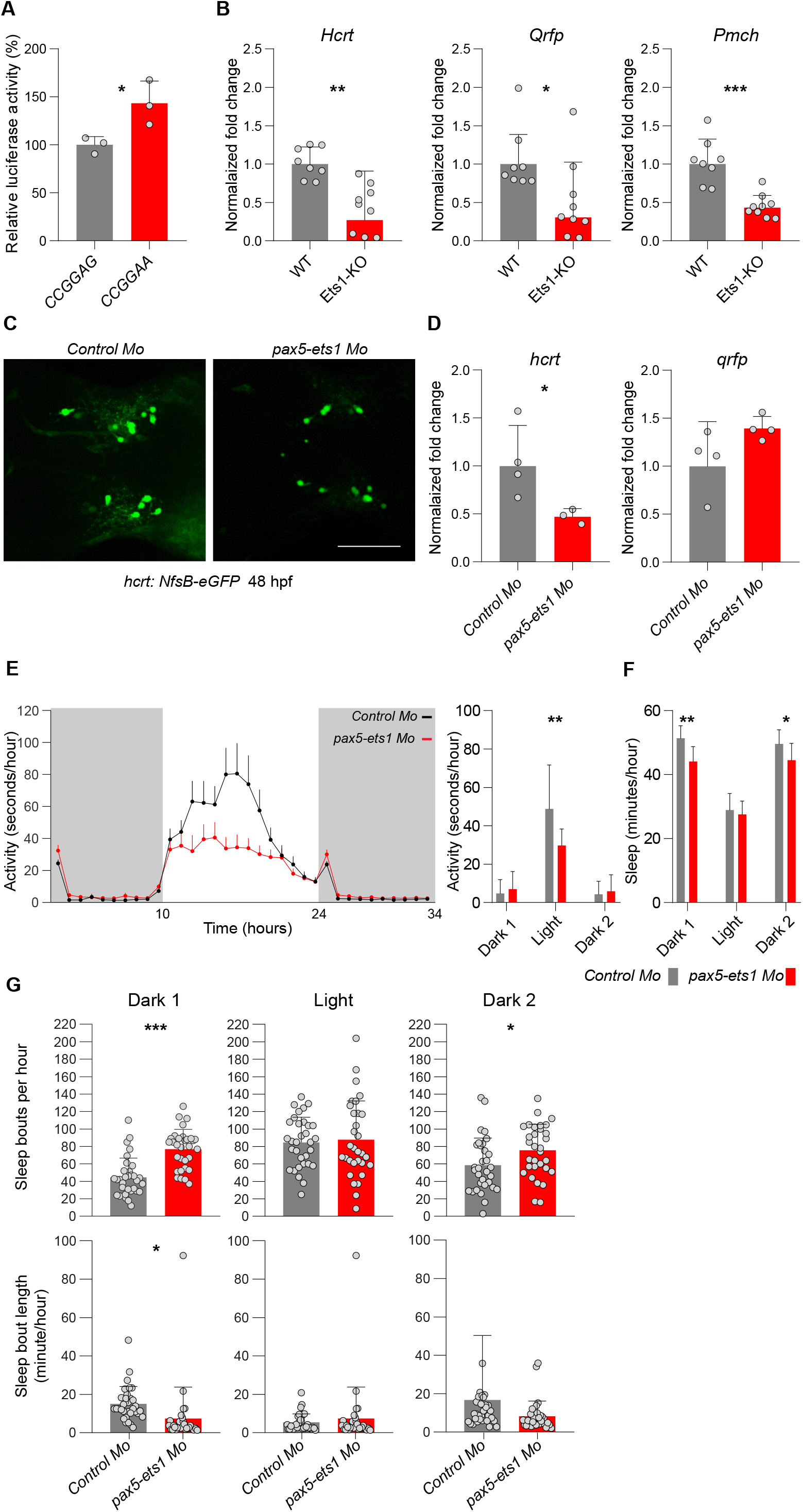
ETS1 and PAX5:ETS1 complex regulate HCRT expression. **a**, The optimized ETS1 binding site (5’-CCGGAA-3’ instead of 5’-CCGGAG-3’) within the human *HCRT* promoter significantly increases the expression of luciferase reporter. * *P* < 0.05, *t*-tests on CCGGAA compared to CCGGAG, N=3. **b**, *Hcrt* expression is significantly decreased in Ets1-KO mice. *Pmch* and *Qrfp* are also significantly decreased, suggesting ETS1 may have a general transcriptional role for lateral hypothalamic neuropeptides. * *P* < 0.05, and *** *P* < 0.001, *t*-tests on log transformed normalized fold change of KO mice (N=9) compared to wild-type controls, N=8 wild-type and N=9 Ets1-KO. Bars represent geometric mean + geometric SD. **c**, Zebrafish larvae injected with morpholinos (Mo) against both *ets1* and *pax5* exhibit a substantial decrease in *hcrt:nfs-eGFP* expression (GFP immunoreactivity 48 hours post fertilization (hpf), scale bar: 50 μm), and hcrt mRNA level (**d**), relative to fish injected with a control morpholino, while the level of expression of *qrfp* is not changed or has tendency to increase. * *P* < 0.05, *t*-test on log transformed normalized fold change of *pax5* and *ets1* morpholinos-injected fish compared to control morpholino-injected fish (N=4 RNA samples extracted from pooled 60 fish at 5dpf each). Knockdown of *ets1* and *pax5* also leads to (**e**) decreased spontaneous locomotor activity during the light period (dark periods are indicated in gray), (**f**) decreased sleep during the dark period, and (**g**) increased sleep fragmentation in the dark period (i.e., a higher number of sleep bouts, of shorter average duration). Recording corresponds to fish at 5-6dpf. N=32, mean+SD; * *P* < 0.05, ** *P* < 0.01, *** *P* < 0.001, *t*-tests.

To assess the significance of the ETS1 binding site for endogenous *Hcrt* gene-expression in live mice, we next quantified *Hcrt* transcripts in the hypothalamus of Ets1-KO mice. A significant decrease in *Hcrt* gene expression was observed as compared to their wild type littermates (Fig. 3b), indicating that ETS1 controls the expression of the mouse *Hcrt* gene *in vivo*, consistently with our *in vitro* data above suggesting it does so for *HCRT*. Additionally, *Qrfp* and *Pmch* expressions were also significantly decreased in Ets1-KO mice (Fig. 3b), suggesting that ETS1 is a major transcription factor controlling the expression of hypothalamic neuropeptides. Interestingly, the methylation of the last 2 CpGs of the *QRFP* promoter was substantially lower than all other CpGs both in the hypothalamus and the blood of narcolepsy patients as well as control subjects (Supplementary Fig. 2). These 2 CpGs belong to a 5’-CGGAAGCCG-3’ sequence, which contains an ETS1 binding site (underlined). This suggests that (i) *QRFP* expression may also be controlled by ETS1, (ii) unlike *HCRT*, the hypomethylated ETS1 binding site of *QRFP* may reflect its occupancy by ETS1 and preserved expression in the hypothalamus of narcolepsy patients.

It was reported that while ETS1 alone has low binding affinity for a suboptimal binding site (5’-CCGGAG-3’), the PAX5 transcription factor is able to recruit ETS1 and forms a PAX5:ETS1 complex which mediates transcription enhancement^31^. To determine whether this complex regulates *hcrt* gene expression, we knocked down both *pax5* and *ets1* genes by injecting morpholino oligonucleotides in zebrafish embryos at the one-cell stage and assessed fish endogenous *hcrt* transcript levels. Fish with normal morphogenetic development (Supplementary Table 2) were selected for gene expression and activity/sleep analysis. Use of a transgenic line expressing a zebrafish *hcrt* promoter-driven eGFP reporter^33^ allowed us to show that while *hcrt* gene expression was downregulated by *pax5-ets*1 knockdown, the number of hcrt-positive cells was unchanged (Fig. 3c). Fish with *pax5* and *ets1* knockdown showed decrease *hcrt* transcript levels, relative to fish injected with a control morpholino oligonucleotide, while the expression of *qrfp* was unchanged (Fig. 3d). Furthermore, rest-activity analysis revealed that *pax5* and *ets1* knockdown fish exhibited a significant decrease in spontaneous locomotor activity during the light period (Fig 3e), while sleep was decreased (Fig 3D) and largely fragmented during the major sleep period (dark) (Fig 3g), a trait commonly seen in narcolepsy^34^. This observation lends further support to the hypothesis that decreased *hcrt* expression, not necessarily cell loss, can cause narcolepsy symptoms. Altogether, our results suggest that ETS1 may have a conserved function in controlling *HCRT* gene expression in man, mouse, and fish.

We propose that the hypothalamus of narcolepsy patients contain HCRT neurons that do not productively express *HCRT*. Epigenetic silencing of HCRT neurons may be mediated by immune mechanisms, for instance involving the release of proinflammatory cytokines within the lateral hypothalamus concurrent to a viral infection. Supporting this view, a polymorphism in the TNF-alpha gene was found to be significantly associated with narcolepsy^35,36^ and narcolepsy patients are reported to have increased serum TNF-alpha levels^37,38^. Moreover, TNF-alpha was found to dramatically decrease *HCRT* and *HCRTR2* expression *in vitro*^39^, and prolonged exposure to TNF-alpha can induce promoter methylation *in vitro*^40^. Intriguingly, a rare condition called autosomal dominant cerebellar ataxia, deafness and narcolepsy, is caused by dominant mutations in the gene encoding the maintenance DNA methyltransferase 1 (*DNMT1*)^41,42^. Such mutations can lead to site-specific hypermethylation and gene silencing^43^, strengthening the notion of a causal link between hypermethylation and narcolepsy. Partial or transient epigenetic modulation of HCRT expression might also be involved in narcolepsy without cataplexy, idiopathic hypersomnia or Kleine-Levin syndrome. Furthermore, the gradual decrease in the number of HCRT-immunoreactive neurons with aging^44^ might also be due to epigenetic downregulation of *HCRT* gene expression, leading to major changes in sleep and sleepiness in aged subjects.

Methylation being reversible, our findings open the avenue for therapeutic interventions based on DNA demethylation, leading to treatment or even cure of narcolepsy. Although DNA methylation/demethylation processes were thought to be restricted to dividing cells, recent evidence indicates that these epigenetic events can occur in post-mitotic cells, including neurons^45^. Novel epigenetic editing strategies, based on both pharmacological reagents or using site-specific demethylating genetic tools, are becoming available^46^. Beyond narcolepsy, epigenetic silencing may represent a key causative factor in other immune, autoimmune and neurodegenerative conditions.

## Methods

### Animals

Hcrt*-*KO and Hcrt-ataxin mice were bred locally on C56BL/6J background. Brains of Hcrt-DTA, Ets1-KO and their controls were provided by international collaborators. Brains of Hcrt-DTA mice were sampled 4 weeks after doxycycline removal from the diet, while their controls were maintained under doxycycline. All animal procedures followed Swiss federal laws and were approved by the State of Vaud Veterinary Office. At all times, care was taken to minimize animal discomfort and avoid pain.

### Human tissue samples

Hypothalamic RNA samples from 4 narcolepsy patients and 4 controls were available in the laboratory of Dr. M. Honda^1^ and were tested locally by qPCR with TaqMan assays Hs01891339_s1 (*HCRT*), Hs01650960_s1 (*QRFP*), and reference genes Hs00744842_sH (*TUBA1B*), Hs01375212_g1 (*RPS18*), and Hs04185005_g1 (*RPS27*). Formalin fixed sections (40μm, 2-4 sections/subject) through the lateral hypothalamus from narcolepsy patients and controls were obtained through international collaborators. All tissues samples were from postmortem brains previously published^2-6^. The research protocol and the use of human biological material was approved by the local ethical committee (SwissEthics). Formalin fixed sections were washed with PBS and genomic DNA was extracted with AllPrep DNA FFPE Tissue kit (Qiagen). CSF and serum samples from narcolepsy and control subjects were provided by the laboratory of Dr. GJ. Lammers.

### RT-qPCR

Mouse brains, cell culture samples and fish were collected and preserved at −80 °C until RNA extraction. Total RNA was isolated using the RNeasy Micro kit (QIAGEN 74004) or Trizol extraction method. The quantity and quality of RNA samples was checked using a Nanodrop (ND-1000) spectrophotometer. Reverse transcription was carried out by using M-MLV reverse transcriptase (Promega) or SuperScript IV reverse transcriptase (Thermo Fisher Scientific) by random hexamer or oligo-dT primers according to the manufacturer’s instructions. Amplification was carried out with TaqMan assay kits Mm01964030_s1 (*Hcrt*), Mm01701538_m1 (*Qrfp*), Mm0124886_g1 (*Pmch*), reference genes Mm01973893_g1 (*Eef1a1*), Mm00850060_s1 (*Rps9*). qPCR for fish samples were performed by primers in combination with Power SYBR Green (Thermo Fisher Scientific) using a ViiA7 real-time PCR system (Thermo Fisher Scientific). Primers were: *hcrt*-F 5’-CTCCTGCAAACTCTACGAGATG-3’, *hcrt*-R 5’-GTCGTTGTTGAGATGCACTAAA-3’, *qrfp*-F 5’-AATGCTGCCACACCAACCAA-3’, *qrfp*-R 5’-TCCCAGGTCCTGAAACAAAGC-3’, and the reference gene *18s*-F 5’-AGCGTGCGGGAAACCACGAG-3’ and *18s*-R 5’-AAGCCGCAGGCTCCACTCCT-3’. Normalized fold expression was calculated according to Taylor *et al*.^7^

### Immunofluorescence and confocal microscopy of human tissues

Hypothalamic formalin-fixed paraffin sections (5μm) were dewaxed in xylene for 15 min, followed by rinsing in 99% and 96% ethanol (3 times, 5 min each). Endogenous peroxidase was blocked for 20 min with methanol / 0.3% H_2_O_2_ followed by rinsing in 96%, 70% ethanol and 1xPBS (3 times, 5 min each). Antigen retrieval was achieved by microwave heating for 3 cycles of 10 sec every 5 min at max 800 W up to 85-9°C in 10mM Citrate Buffer (pH 6.0), then cooling down to room temperature in the same buffer for 1 hour. Slides were washed in 1x PBS (3 times, 5 min each) and incubated in bovine serum albumin (2% BSA, 0.3% Triton x-100, 5% normal donkey serum in 1xPBS) for 30 min. QRFP antibody^8^ at 1:400 was applied overnight at room temperature, followed by a second incubation with QRFP antibody 1:400 and HCRT antibody 1:500 (goat HCRT expression (yellow arrows) anti HCRT, Santa Cruz Biotechnology, SC-8070) at 4 °C for 24 hours. After rinsing with 1xPBS for 3 times (10 min each), secondary antibodies were applied (donkey IgGs coupled to Alexa-594, or -488 fluorophores) ins 1:500 dilutions for 2 h at room temperature followed by rinsing with 1xPBS for 3 times (10 min each). To test the specificity of QRFP antibody, QRFP antibody was blocked with QRF (NM_198180) Human Over-expression Lysate (2μg/ml, 1%BSA) for 2 hours at room temperature before use in the above described immunofluorescence assay. Images were acquired on an inverted Zeiss LSM780 confocal laser-scanning microscope (405, 488, and 561 nm lasers) using a 40x oil objective (EC plan-Neofluar 40x/1.30 Oil DIC M 27). For each human section used for cell quantification, confocal images covering the HCRT positive cell field were acquired at 8-bit image depth and a frame of 1,024 × 1,024 pixels, and tiled together using ZEN software at size of 2125.48×2125.48 μm. Immunoreactive cell counts were evaluated within that frame using ImageJ software.

### QRFP radioimmunoassay

Quantification of QRFP in human CSF samples was carried out using a specific radioimmunoassay (RIA), previously described in detail^8^. CSF samples were pumped at a flow rate of 1.5 ml/min through one Sep-Pak C_18_ cartridge. Bound material was eluted with acetonitrile/water/TFA (50:49.9:0.1; v/v/v) and acetonitrile was evaporated under reduced pressure. Finally, the dried extracts were resuspended in PBS 0.1 M and assayed for QRFP. Data represent the mean of duplicates for each sample.

### *HCRT* promotor cloning

1800 base pairs of the human *HCRT* promoter upstream of the transcription start site, were cloned in multiple cloning site of the pCpG-basic vector^9^ upstream of Firefly luciferase sequence. The cloned plasmids were transformed in a Pir1 bacterial strain and sequenced for verification. The bacterial colonies were cultured with Zeocin selection at large scale and maxi-plasmid preparations were carried out using HiSpeed Plasmid Maxi Kit (Qiagen, 12662). CpG Methyltransferase, M.SssI (New England Biolabs, M0226s) was used to methylate the CpG sites on the *HCRT* promoter.

### Luciferase assay

HEK 293FT and PC12 cell lines were cultured at 1.5-2*10^5^ density in 24-well plates in DMEM medium containing 10% FBS. Cultures were transfected with methylated and unmethylated plasmids. The transfection was performed using X-tremeGENE™ HP DNA Transfection Reagent (Sigma, 6366244001) with the DNA to reagent ration of 1:3. The transfected plasmids were mixed with Firefly and Renilla luciferases with ratio of 19:1. 48 hours after transfection the cells were lysed and prepared for the luciferase activity measurement according to Promega kit (E1910). Luciferase activity was measured using GloMax 96 microplate luminometer from Promega. The Firefly luciferase activity was normalized to the Renilla activity.

### DNA methylation analysis

Five hundred ng of DNA was treated with sodium bisulfite using the EpiTect Fast DNA Bisulfite kit (Qiagen). Primers for nested PCR (2 fragments for *HCRT*, 1 for *PDYN and HCRTR2*) were designed with MethPrimer^10^ (Supplementary Table 1).

Two methods were used to quantify CpG methylation. In the first method PCR amplicons were gel purified followed by Sanger sequencing. Sequence trace files were analyzed by ESME software^11^ resulting in average methylation at each CpG. In the second method purified PCR amplicons (100 ng) were used to prepare sequencing libraries by TruSeq Nano DNA LT Library Preparation Kit (Illumina) with the following modifications. Purified bisulfite PCR products were not sheared, but directly subjected to end repair and then cleaned up with a single round of purification beads at a 1.8X ratio. The standard protocol was strictly followed for the subsequent steps. Libraries were quantified with a fluorimetric method (QubIT, Thermo Fisher) and their size pattern evaluated with the Fragment Analyzer (Agilent). An equimolar pool of each librarie was prepared and loaded at 10 pM (with 25% PhiX spike in for generating sequencing diversity) for a 150 cycle paired-end run on a MiSeq instrument (Illumina) using a v2 reagent kit. Sequencing data were demultiplexed using the bcl2fastq conversion software (version 2.20, Illumina). Quality control was performed using the NGmerge tool^12^ to keep only read-pairs that could be reconstructed (between 59 and 67% of the reads). On average, 160194 reads (range 10824-349712 reads) were obtained par fragment and per subject. A weight matrix profile was prepared from the reference sequence to take into account the C to T alterations caused by bisulfite treatment and the reads were aligned to the profile using pfsearch from pftools^13^. A summary table counting the number of occurrences of each nucleotide along the fragment of interest was produced and the ratio of methylated C nucleotides was computed as a percentage of methylation for each position.

### Morpholino study in zebrafish

Morpholino antisense oligos targeting *ets1* and *pax5* transcripts were designed according to Pham *et al*.,^14^ and Kwak *et al*.,^15^, respectively. 0.25pmol of *ets1* (5’-GTCATGGTCACGCATTCAAACGTAC-3’) or a combination of 0.375pmol of *pax5* TB1 (5’-CAGTGGATTTCCATCTGTTTTAAA-3’) and 0.375pmol of *pax5* TB2 (5’-CTCGGATCTCCCAGGCAACATGGT-3’) were injected in 1-2 cells zygote in a total volume of 1nl. The control morpholino corresponds to the standard sequence proposed by the manufacturer (5’-CCTCTTACCTCAGTTACAATTTATA-3’). Morpholino synthesis was performed by Gene-tools (http://www.gene-tools.com).

Visualization of Hcrt neurons was performed using the *Tg(hcrt:nfsB-EGFP*) transgenic line, in which a 2-kb fragment of the zebrafish hcrt promoter drives expression of a nfsB-EGFP fusion protein^16^. Imaging was performed on homozygous embryos, using a LSM710 confocal microscope (Zeiss). qPCR and locomotion tracking were performed on wild-type AB line. Embryos and larvae were maintained in an incubator at 28.5°C on 14/10h light/dark cycle and staged as described by Kimmel *et al*.,^17^. Rest-activity quantification was performed according to Seifinejad *et al*.,^18^. Briefly, larvae were recorded from 5 to 7dpf in 96-well plates, one fish per well, in 300μl of egg water. Individual activity was measured with the quantification mode of Zebrabox (viewpoint) set with the following parameters: Freeze: 3, Burst: 29, Threshold: 15, Bin: 1min, Light intensity: 7%. Any 1min with less than 0.1s of movement was counted as 1min of sleep. Larvae exhibiting morphological defects (table S2) were excluded from analysis to control against motor impairments.

### Statistics

All statistics were performed using GraphPad Prism 8 and are reported in figure legends.

## Data availability

All data are available in the main text and the supplementary materials.

## Acknowledgments

We thank T. Scammell, D. Swaab, Y. Dauvilliers, and C. Peyron for providing human brain tissues. We thank the Lausanne Genomic Technologies Facility for help in NGS-methylation sequencing. We thank T. Kilduff and A. Yamanake for providing orexin-DTA mice brains, H. Okamoto for providing *Tg(hcrt:nfsB-eGFP)* fish, and M. Rehli for providing the CpG-free vector. We thank also T. Miyagawa and M. Shimada from Tokyo Metropolitan Institute of Medical Science for assistance. We thank all the persons at the zebrafish facility of the University of Lausanne. We are grateful to U. Schibler and R. Jaenisch for comments on our draft manuscript. This work was supported by E-RARE NARCOMICS (Swiss National Science Foundation grant 185655 to M.T.), Swiss National Science Foundation (grant 173126 to M.T.), and the State of Vaud (Faculty of Biology and Medicine, University of Lausanne). The zebrafish work was supported by the Swiss National Science Foundation (grant 188789 to F.A.). L.S. has received funding from the European Union’s Horizon 2020 research and innovation program under the Marie Skłodowska-Curie grant (Agreement No. 707404).

## Author contributions

A.S., A.N., R.L., F.A., A.V., G.J.L., and M.T. conceived the project. A.S., A.N., S.L., C.F., R.F., L.S, L.A.G-S., D.F., M.H., Y.A., D.G., M.P., N.C., and M.T. carried out experiments and analyzed data. C.I. contributed to NGS sequence analyses. A.S., A.N., R.L., A.V. and M.T. wrote the paper with contribution from all authors.

## Additional information

**Supplementary Fig. 1.**
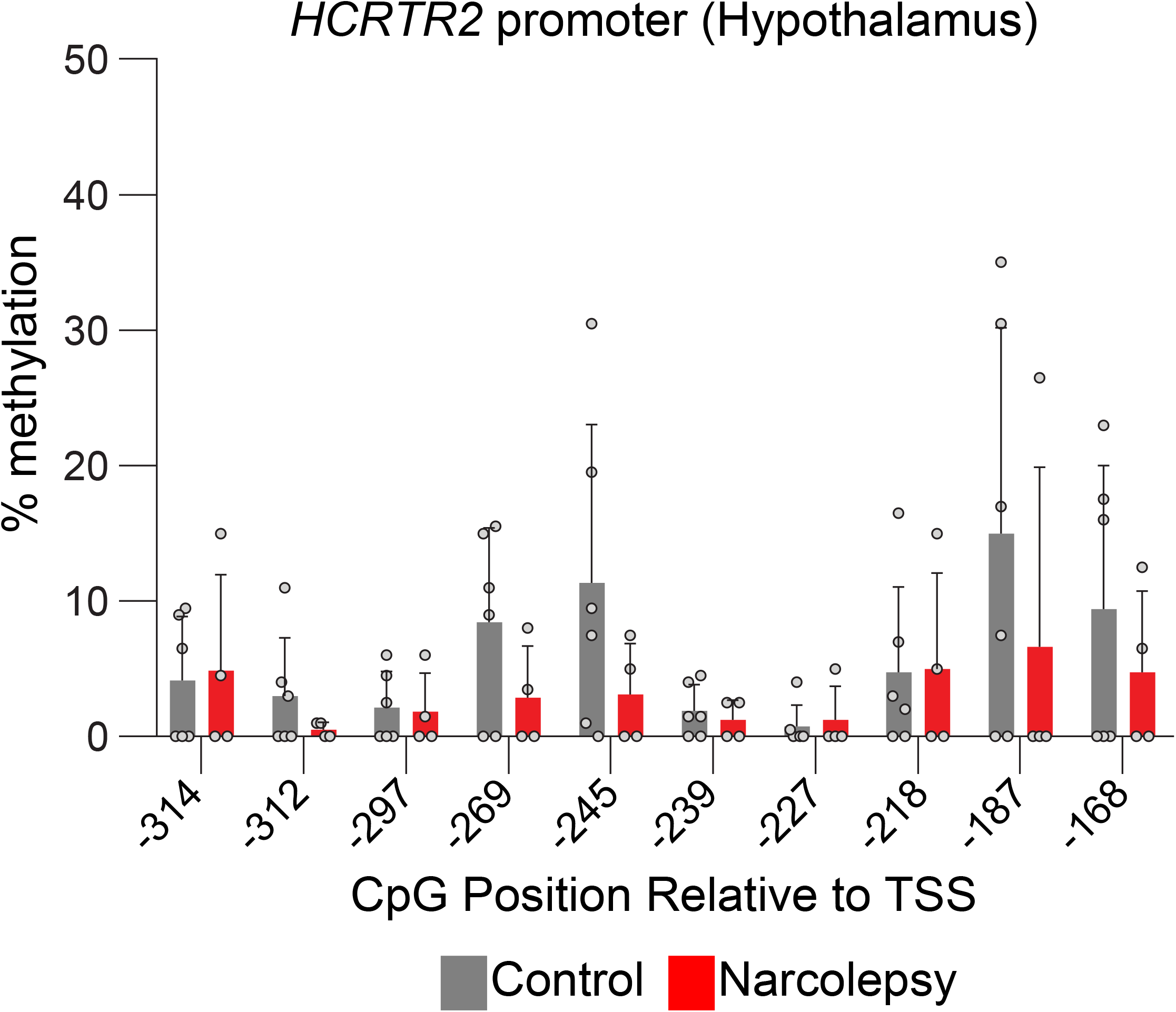
Methylation status of the human *HCRTR2* promoter in the hypothalamus of narcolepsy patients and controls.

**Supplementary Fig. 2.**
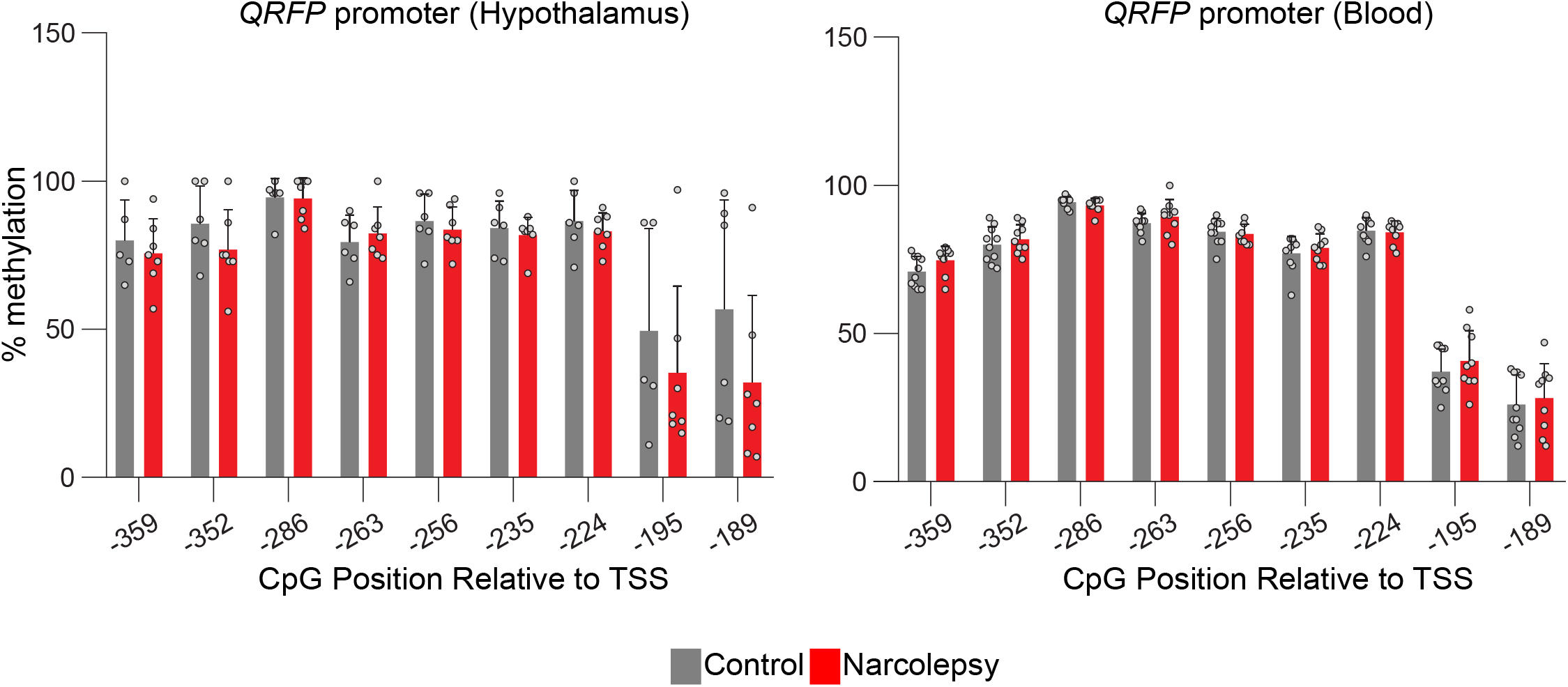
Methylation status of the human *Qrfp* promoter in the hypothalamus and blood of narcolepsy patients and controls.

**Supplementary Fig. 3.** Representative fish image showing normal phenotype after Mo injection.

**Supplementary Table 1:** Primers used for bisulfite sequencing.

**Supplementary Table 2:** Number and percentage of zebrafish without developmental abnormalities used in behavioral assays.

**Table S1.**
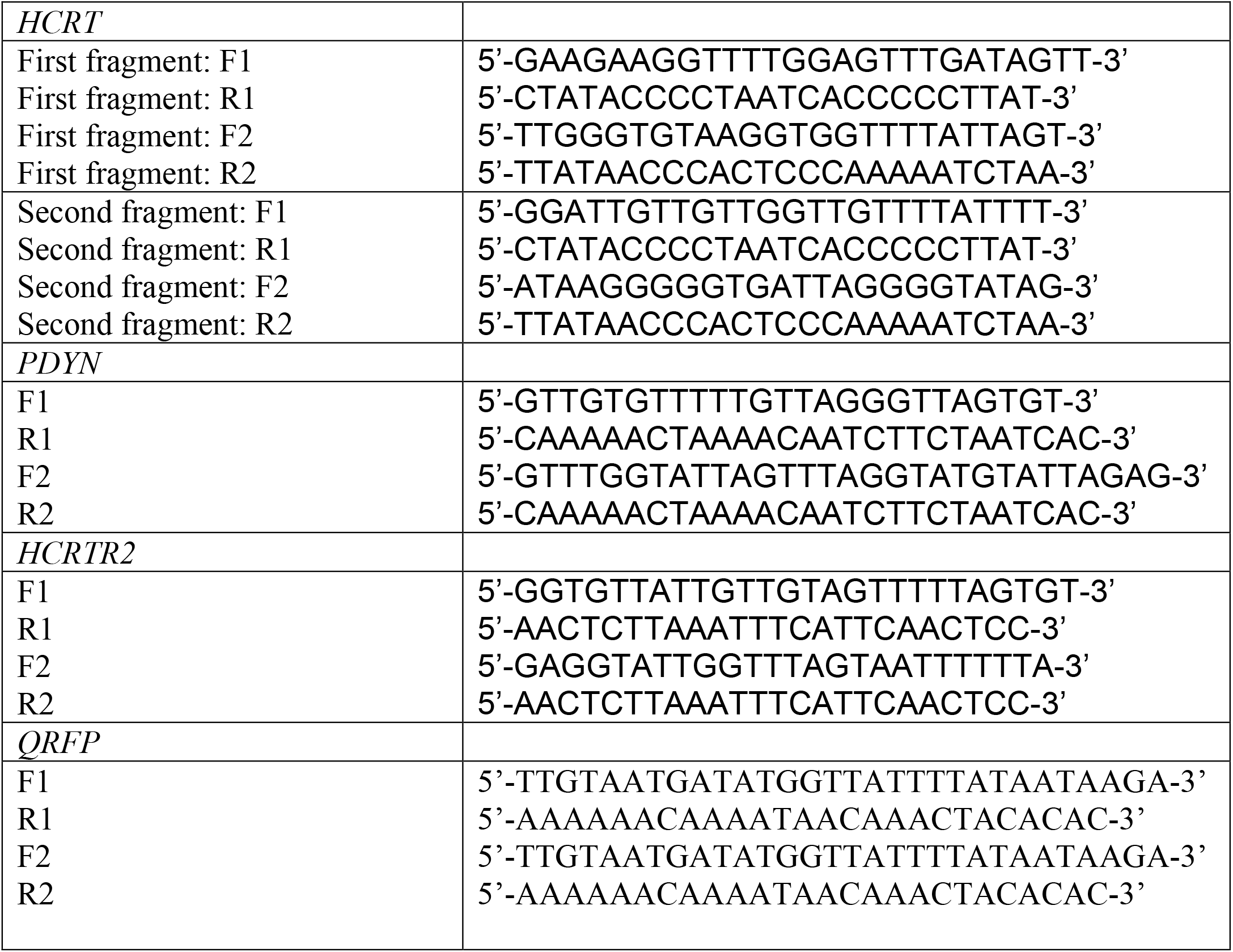
Nested bisulfite primers used in this study.

**Table S2:** Number and percentage of zebrafish without developmental abnormalities used in behavioral assays.

|  | Injection 1 |  | Injection 2 |  | Injection 3 |  | Injection 4 |  |
| --- | --- | --- | --- | --- | --- | --- | --- | --- |
|  | Ctrl Mo | Mo Pax5-Ets1 | Ctrl Mo | Mo Pax5-Ets1 | Ctrl Mo | Mo Pax5-Ets1 | Ctrl Mo | Mo Pax5-Ets1 |
| Normal Morphology | 70 | 36 | 86 | 82 | 37 | 42 | 56 | 78 |
| Deformities | 0 | 7 | 1 | 10 | 1 | 5 | 0 | 12 |
| % Deformities | 0.00% | 16.28% | 1.15% | 10.87% | 2.63% | 10.64% | 0.00% | 13.33% |

